# Loss of rice *PARAQUAT TOLERANCE 3* confers enhanced resistance to abiotic stresses and increases grain yield in field

**DOI:** 10.1101/2020.02.22.961151

**Authors:** Alamin Alfatih, Jiu Wu, Sami Ullah Jan, Zi-Sheng Zhang, Jing-Qiu Xia, Cheng-Bin Xiang

## Abstract

Plants frequently suffer from environmental stresses in nature and have evolved sophisticated and efficient mechanisms to cope with the stresses. To balance between growth and stress response, plants are equipped with efficient means to switch off the activated stress responses when stresses diminish. We previously revealed such an off-switch mechanism conferred by Arabidopsis *PQT3*, knockout of which significantly enhances resistance to abiotic stresses. To explore whether the rice homolog *OsPQT3* is functionally conserved, we generated three knockout mutants with CRISPR-Cas9 technology. The *OsPQT3* knockout mutants (*ospqt3*) display enhanced resistance to oxidative and salt stress with elevated expression of *OsGPX1*, *OsAPX1*, and *OsSOD1*. More importantly, the *ospqt3* mutants show significantly enhanced agronomic performance with higher yield compared with the wild type under salt stress in greenhouse as well as in field conditions. We further showed that *OsPQT3* was rapidly down regulated in response to oxidative and other abiotic stresses as *AtPQT3*. Taken together, these results support our previous findings that AtPQT3 acts as an off-switch in stress response, which is well conserved in rice. Therefore, *PQT3* locus provides a promising candidate for crop improvement with enhanced stress resistance by gene editing technology.

## INTRODUCTION

Oxidative stress is a common component to all environmental stresses that disturb cellular oxygen metabolism (Das and Roychoudhury, 2014). Plants have evolved well-coordinated defense mechanisms to cope with oxidative stress generated under wide-range abiotic stresses (Mittler et al., 2004). Two eminent types of defensive responses are enzymatic response and non-enzymatic response. The first type of response comprises of cellular enzymes like superoxide dismutase (Li et al., 2011), peroxidase (POD), catalase (CAT), ascorbate peroxidase (APX), peroxiredoxin Q (PRXQ), and glutathione peroxidase (GPX) which protect cell by engaging toxic ions entering the cell or generated within the cell in stress conditions (Apel and Hirt, 2004; Mittler et al., 2004; Takahashi and Asada, 1988). Such antioxidant enzymes are helpful in achieving tolerance against oxidative stress (Ye and Gressel, 2000). The later type of defense response include non-enzymatic cellular components such as ascorbate and glutathione that, in majority, serves as antioxidants as well as signals to modulate cellular functions in response to abiotic or biotic stimuli (Moon et al., 2004; Vierstra, 2009).

Paraquat (*N,N’-dimethyl-4,4’-bipyridinium dichloride*, PQ), a widely used non-selective and rapid-acting herbicide, is a highly effective inducer of oxidative stress in plants. Paraquat diverts electrons from photosystem I towards molecular oxygen, generating superoxide, which is rapidly converted by superoxide dismutase to hydrogen peroxide, and thus raising the concentration of reactive oxygen species (ROS) to toxic or even lethal level in green plants (Summers, 1980), which alters normal cellular homeostatic condition (Haley, 1979; Miller et al., 2010). The accumulated ROS damages macromolecules and biological membranes, induces cell senescence, and causes irreversible injuries to cell and cellular components (Gill and Tuteja, 2010). It is well established that ROS concentration significantly and rapidly increases under numerous stresses such as drought, salt and paraquat-induced stress (Leshem et al., 2006; Miller et al., 2010; Van Breusegem et al., 2008).

Paraquat is also a commonly used oxidative stress inducer in the laboratory for mutant screen (Chen et al., 2009; Kurepa et al., 1998). We previously isolated paraquat resistance mutants (Xi et al., 2012) and reported *AtPQT3* encoding an E3 ubiquitin ligase, which turns off or negatively regulates the response activated against oxidative stress (Luo et al., 2016). AtPQT3 interacts with PRMT4b that help plants tolerate oxidative stress by methylating histones on the chromatin of *AtGPX1* and *AtAPX1*, thus enhancing their expression and leading to stress resistance (Luo et al., 2016). In the process of ubiquitination, E3 ubiquitin ligases regulate downstream signaling processes in biotic and abiotic stress response through mediating transcription factors (TFs) or vesicle trafficking (Zhou et al., 2017). An example reported earlier showed that E3 ubiquitin ligase RING-type MIEL1 interacts with AtMYB30 leading to proteasomal degradation of *AtMYB30,* thus causing *Arabidopsis* defense attenuation (Marino et al., 2013). Another RING-type E3 ubiquitin ligase EIRP1 interacts with WRKY11 in *Vitis pseudoreticulata* promoting defense against fungal infection (Yu et al., 2013). Similarly, *OsHCI1* is responsive to temperature variation in rice. When overexpressed, *OsHCI1* showed improved tolerance to heat in *Arabidopsis* (Lim et al., 2013). Lately, it was reported that the E3 ubiquitin ligase *SOR1* modulates ethylene response by regulating IAA/Aux protein stability in rice roots (Chen et al., 2018).

In this study, we investigate whether *OsPQT3*, the rice homolog of *AtPQT3*, is functionally conserved or not. Our results show that *OsPQT3* negatively regulates oxidative stress response. Loss-of-*OsPQT3* confers salt and paraquat tolerance as demonstrated by the *ospqt3* knockout mutants. The expression of *OsAPX1*, *OsGPX1* and *OsSOD1* was increased in *ospqt3* as compared to that in the wild type. Our findings suggest that *OsPQT3* shares functional similarity with *AtPQT3* as both are negative regulators of oxidative stress response. Moreover, *ospqt3* mutants exhibited significantly enhanced growth and grain yield of rice under both normal and stressed conditions, therefore providing a candidate gene for improving abiotic stress tolerance and yield of crops via gene editing.

## RESULTS

### Loss-of-*OsPQT3* confers enhanced tolerance to paraquat and salt

Previous studies in dicots have demonstrated that the *atpqt3* plays important roles in the response of *Arabidopsis thaliana* to oxidative stress (Luo et al., 2016). However, its roles in rice remain unknown. We resorted to CRISPR-Cas9 technology to generate mutations in *OsPQT3* gene and obtained 3 independent mutants (*ospqt3-1, ospqt3-2,* and *ospqt3-3*). The first mutant (*ospqt3-1*) had a 5-base deletion in the first exon, the second mutant (*ospqt3-2*) gained one base in the first exon, and the third mutant (*ospqt3-3*) gained one base in the second exon. Consequently, the open reading frame of *Os10g29560* was interrupted in all three mutants as confirmed by genomic DNA sequencing analyses (Figure 1). Therefore, we successfully generated three independent null mutants for further study.

**Figure 1.**
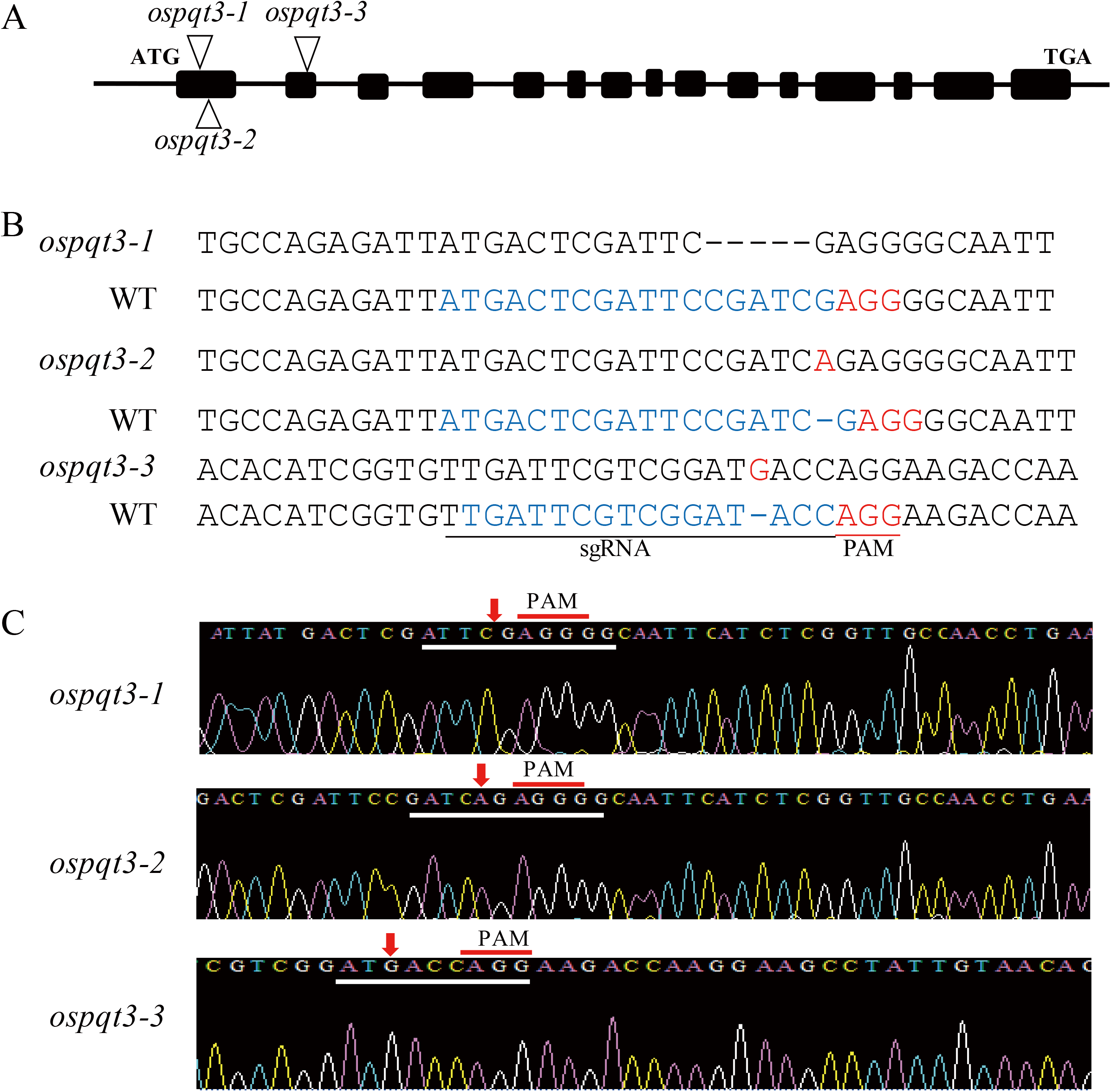
Identification of *pqt3* mutant plants. A. Schematic representation of *OsPQT3* gene and location of CRISPR-Cas9 edited sites (indicated by triangles). B. Sequence around CRISPR-Cas9 edited sites. C. Sequencing confirmation of the edited site in the mutants.

We first checked the germination rate in response to paraquat and salt stress. The seeds of *ospqt3-1*, *ospqt3-2, ospqt3-3* and wild type were germinated in MS medium with or without 0.1 μM paraquat for eight days. The germination of wild type and mutants revealed no difference in the absence of paraquat. However, in the presence of 0.1 μM paraquat the germination of *ospqt3* mutants were significantly higher than the wild type plants (Figure 2A). Similarly, the germination rate of wild type plants were severely decreased in the medium containing 175 mM NaCl compared to *ospqt3* mutant (Figure 2A). This result suggests that *OsPQT3* play a negative role in seed germination in response to abiotic stresses.

**Figure 2.**
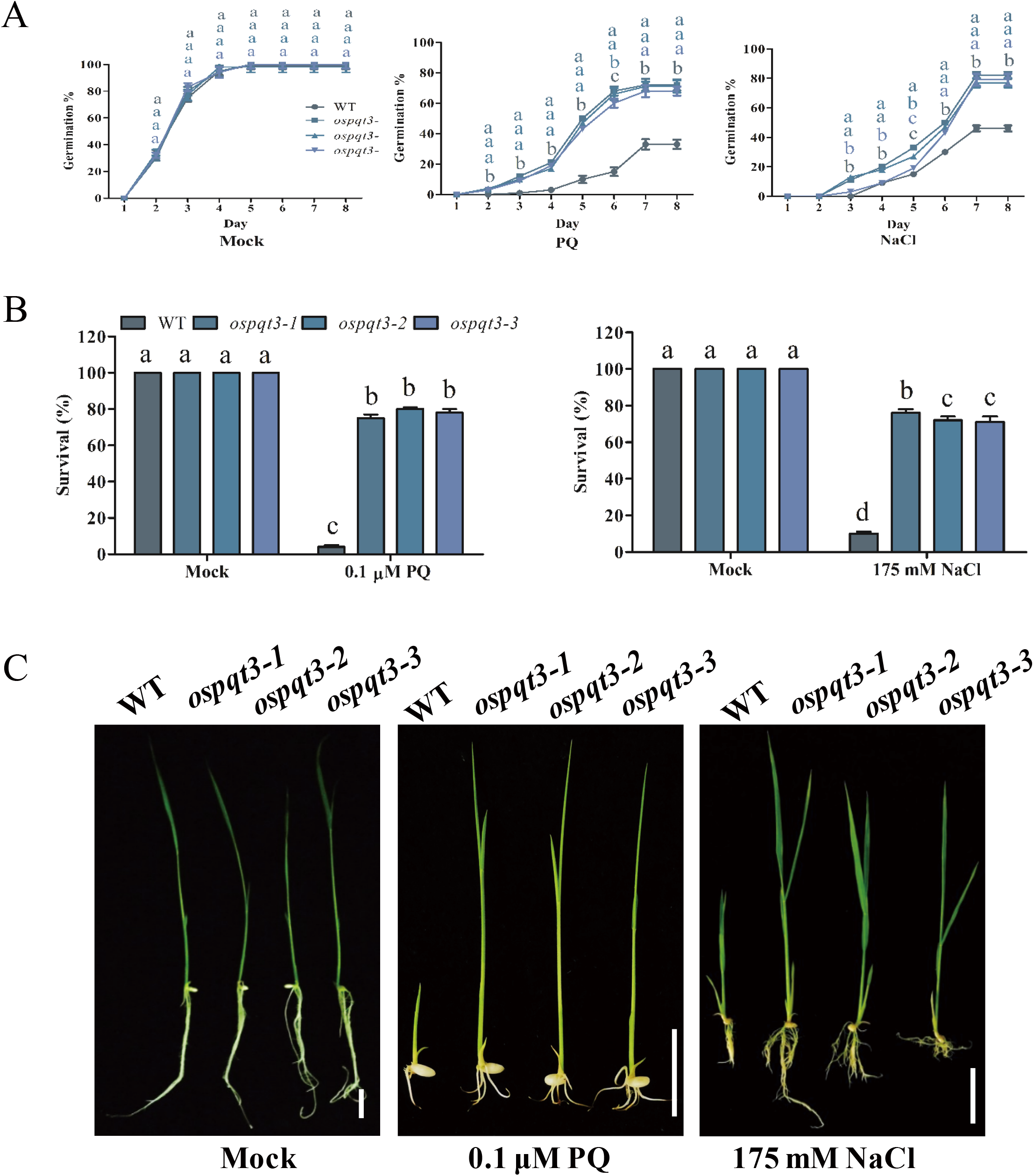
Loss-of-OsPQT3 confers paraquat and salt tolerance phenotypes in seed germination. A. Seed germination curve. Seeds of WT and *pqt3* mutants were germianted on MS medium supplemented with 0 (Mock) or 0.1 μM paraquat (PQ) and 175 mM NaCl for 8 days and germination % was recorded everyday. Values are mean ±SD (20 seeds per replicate, 4 replicates per treatment). Different letters denote significant differences (P < 0.05) from Duncan’s multiple range tests. B. Seedling survival ratio. The above germinated seeds were grown for 2 weeks before seedling survival ratio was recorded. Values are mean ± SD (20 plants per replicate, 4 replicates per treatment). Different letters denote significant differences (P < 0.05) from Duncan’s multiple range tests. C. Growth of germinated seeds. Bar = 2 cm.

Next, we checked the survival ratio of seedlings in response to paraquat and salt stress. The seeds of *ospqt3-1*, *ospqt3-2, ospqt3-3* and wild type were germinated in MS medium with or without 0.1 μM paraquat for two weeks. All the wild type and mutants survived on MS medium without apparent differences. However in the presence of paraquat, the survival ratio *ospqt3* mutants was more than 80% in contrast to less than 20% of the wild type (Figure 2B). This result suggests that the mutants are more tolerant to the oxidative stress generated by paraquat in comparison to wild type. When germinated in presence of 175 mM NaCl for 2 weeks, the survival ratio of the *ospqt3* mutants was around 80%, while the wild type had only 20 % survival rate (Figure 2B). This result indicates that *OsPQT3* plays negative roles in salt stress tolerance in rice, which is consistent to the *Arabidopsis pqt3* mutant (Luo et al., 2016). Furthermore, the growth of the survived wild type seedlings was significantly inhibited in the presence of paraquat and salt compared with the mutants (Figure 2C).

### The *ospqt3* mutants show improved salt stress resistance in the seedling and vegetative stages

To evaluate the attributes of salt resistance in the *ospqt3* mutant seedlings, we treated six-day-old seedlings of *ospqt3-1, ospqt3-2, and ospqt3-2,* and wild type (WT) with 0, 75, 100 mM NaCl in hydroponic solution for 10 days. Both *ospqt3* mutants and WT seedlings grew well under 0 mM NaCl although the wild type shoot was a little taller than the mutant. After exposure to 75 mM or 100 mM salt stress, most *ospqt3* mutants seedlings remained green and showed continuous growth, whereas WT seedlings at 75 mM NaCl were 20 % dead while at 100 mM NaCl were almost 90 % dead (Figure 3A and B). The survival rate of the WT seedlings were 10 %, but the survival rate of *ospqt3* mutant seedlings reached 80% under 100 mM NaCl treatment (Figure 3B).

**Figure 3.**
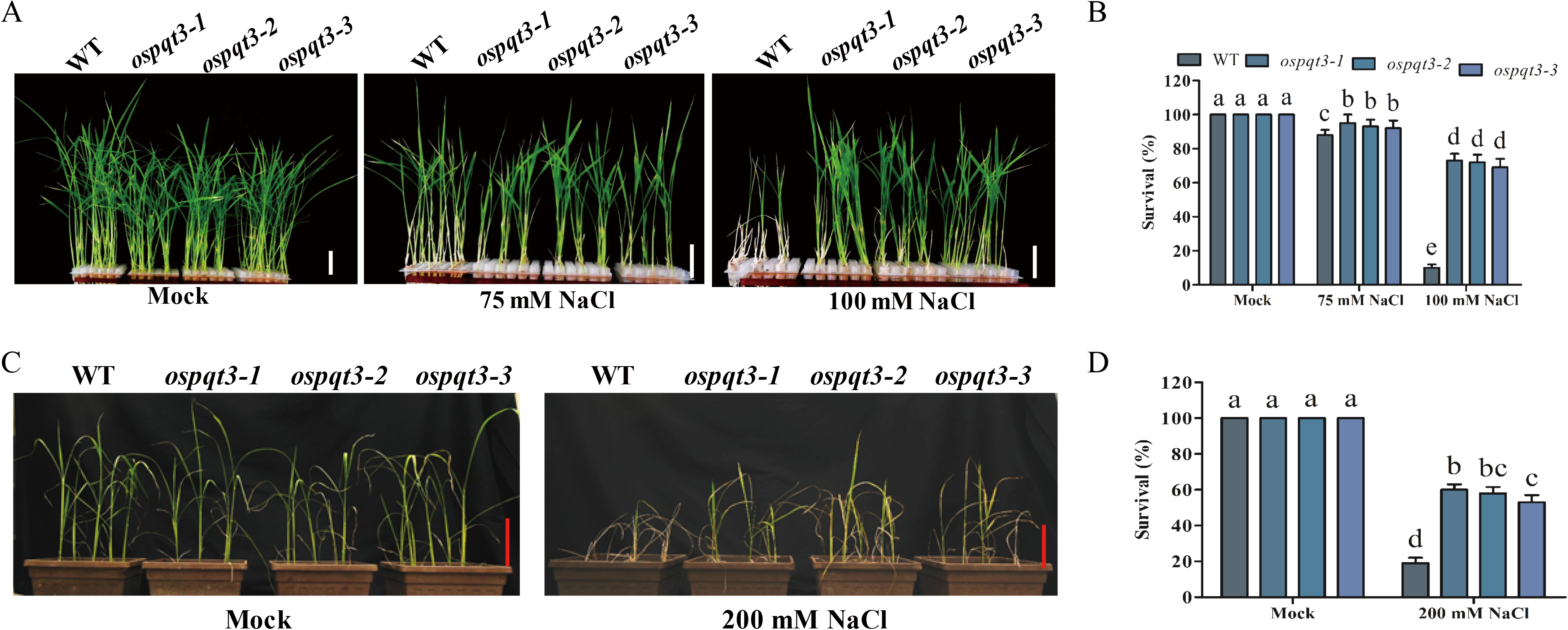
Growth phenotype of *ospqt3-1*, *ospqt3-2* and *ospqt3-3* mutants in the presence of salt. A. Seeds of WT and *pqt3* mutants were germinated and tranferred to hydroponic culture for three days and then the medium was supplemented with 0 mM (Mock), 75 mM or 100 mM NaCl and grown for 10 days before photographs were taken. Bar = 2 cm. B. Survival ratio of the above genotypes and treatments. Values are mean ± SD (24 plants per treatment, 3 replicates per treatment). Different letters denote significant differences (P < 0.05) from Duncan’s multiple range tests. C. Salt tolerance assay in soil. The 4-week-old WT and *ospqt3* mutants plants were watered by 0 mM (Mock) or 200 mM NaCl for 5 days before photographs were taken. Bar = 10 cm. D. Survival ratio of the above genotypes and treament. Values are mean ±SD (5 plants per treatment, 6 replicates per treatment). Different letters denote significant differences (P < 0.05) from Duncan’s multiple range tests.

To evaluate salt tolerance of *ospqt3* mutants grown in soil, we grew plants in the greenhouse and irrigated 4-week-old plants of WT, *ospqt3-1, ospqt3-2,* and *ospqt3-3* with 0 mM or 200 mM NaCl for 1 week (Figure 3C). The result shows that, the mutant plants were significantly more tolerant to the salt stress compared with the WT plants. The survival rate of the *ospqt3* mutants was more than 50 % while 21% of WT survived under 200 mM NaCl treatment (Figure 3D). Collectively, these results suggest that *ospqt3* mutants are more resistant to salt stress than the WT at the seedling and vegetative growth stages.

### The *ospqt3* mutants exhibit improved grain yield in field conditions

To test the field performance of *ospqt3* mutants, we conducted field trials of WT (ZH11), *ospqt3-1, ospqt3-2,* and *ospqt3-3* mutant plants at two different locations in China: Hefei, Anhui Province from April to August in 2019 (Figure 4) and Hainan Island from November to April in 2019 (Figure 5). Figure 4A shows the typical plant of WT and *ospqt3* mutants from the normal field trial in Hefei and Figure 4B shows the panicle size of WT and *ospqt3* mutant plants. The *ospqt3* mutants exhibited significantly increased grain yield by 11.4% (Figure 4C), tiller number by 25% (Figure 4D), panicle length by 12.2% (Figure 4E), seed setting rate by 6.7% (Figure 4F), and primary branch number by 14.2% (Figure 4G), except for plant height (Figure 4H). The field trial results in Hainan Island is shown in Figure 5. A typical panicle is shown for different genotypes in Figure 5A. Again, the *ospqt3* mutants exhibited significantly increased grain yield by 15% (Figure 5B), tiller number by 20% (Figure 5C), and panicle length by 12.5% (Figure 5D) but no difference was found in plant height (Figure 5E). Taken together, these results suggest that *ospqt3* mutants exhibit not only paraquat and salt tolerance, but also enhanced grain yield, making it a promising candidate for crop improvement.

**Figure 4.**
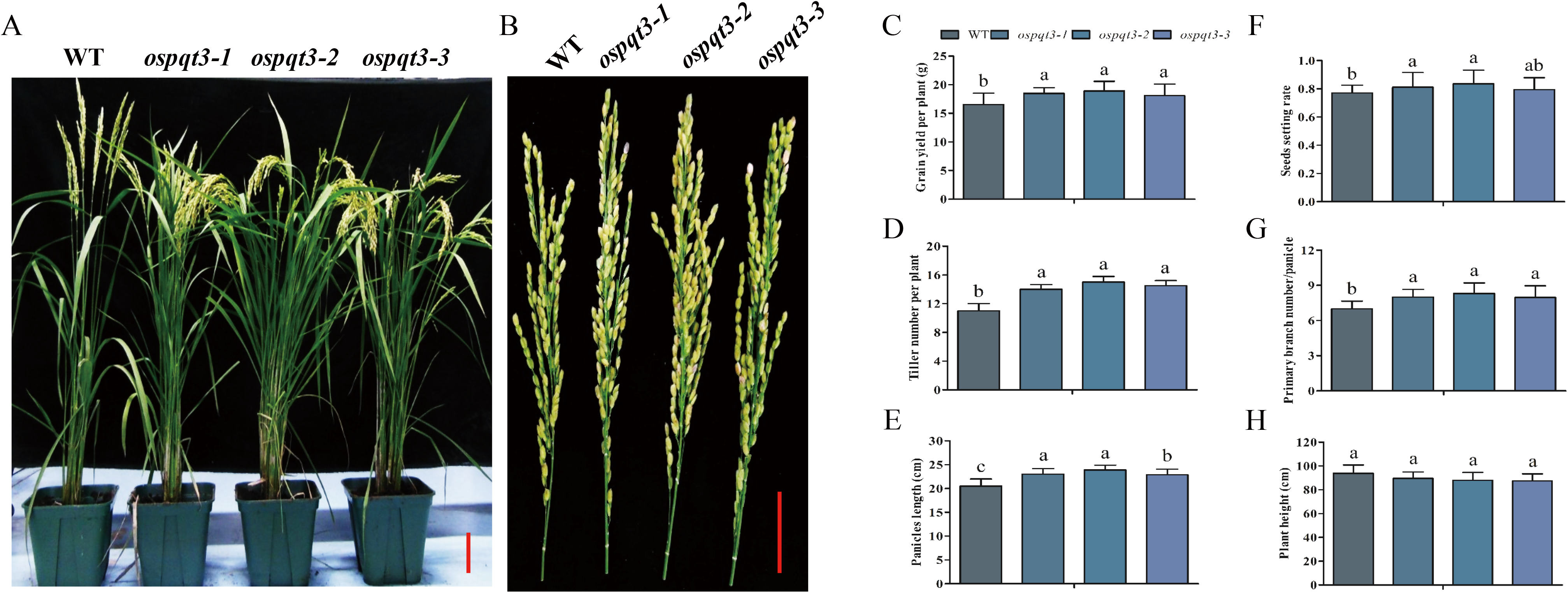
The *ospqt3* mutants exhibit improved yield in the field. A. Growth status of WT (ZH11), *ospqt3-1*, *ospqt3-2*, and *ospqt3-3* mutants plants in the field (Hefei, 2019). Bar = 10 cm. B. Representative panicles of the *ospqt3* mutants and WT (ZH11). Bar = 4 cm. C-H. Yield components and plant height. Grain yield per single plant, tillers number per plant, panicles length, seeds setting rate, primary branch number per panicle, and plant height, respectively. Values are the mean ± SD (30 plants per treatment, 3 replicates per treatment). Different letters denote significant differences (P < 0.05) from Duncan’s multiple range tests.

**Figure 5.**
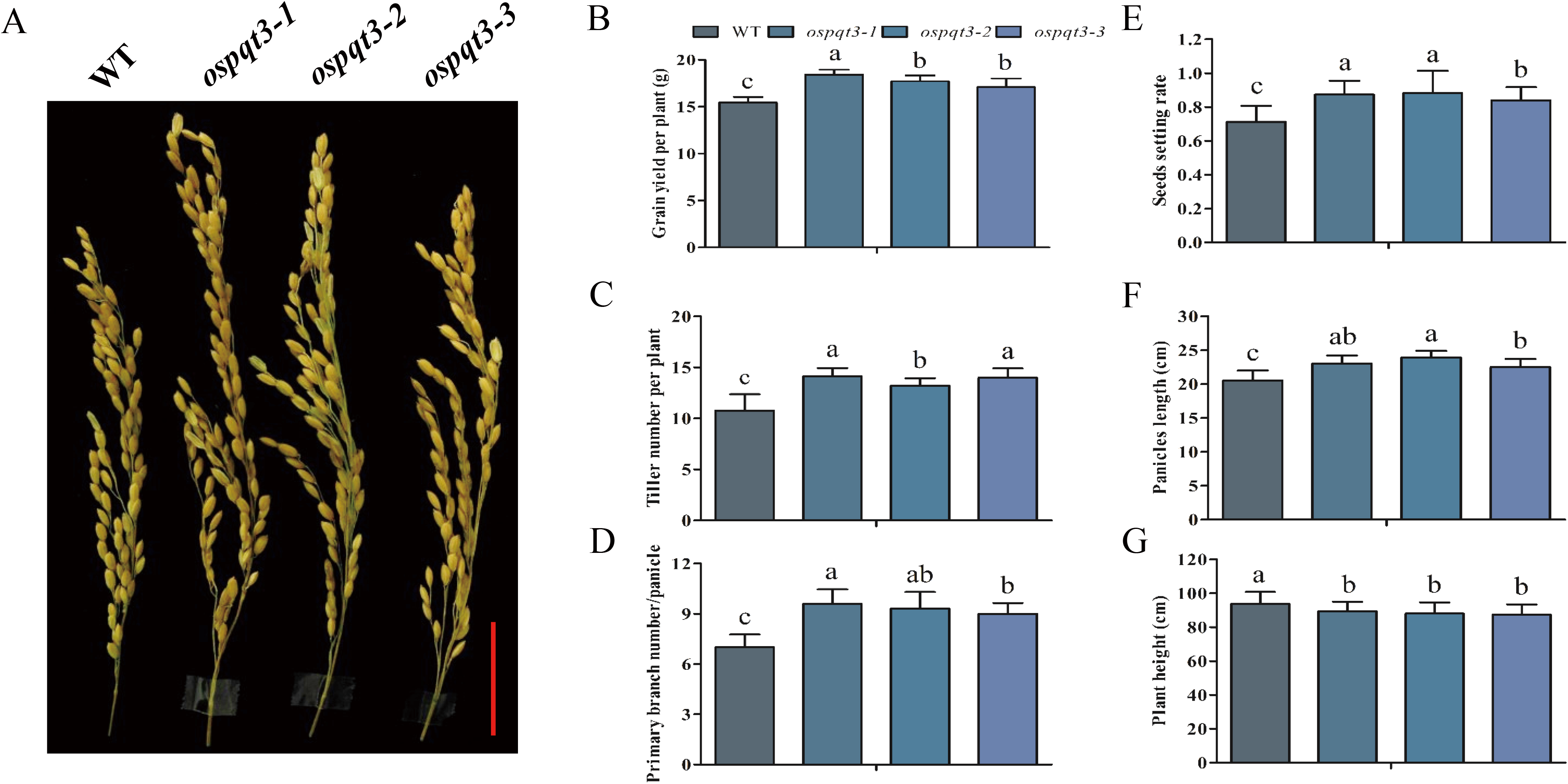
The repeat field trial confirmed the improved yield of *ospqt3* mutants. A. Representative panicles of the *ospqt3* mutant and WT (ZH11) plants grown in the filed (Hainan Island, 2019). Bar = 4 cm. B-G. Yield components and plant height. Grain yield per plant, tillers number per plant, panicles length, seeds setting rate, primary branch number per panicle, and plant height, respectively. Values are the mean ± SD (30 plants per treatment, 3 replicates per treatment). Different letters denote significant differences (P < 0.05) from Duncan’s multiple range tests.

### The *ospqt3* mutants exhibit enhanced salt tolerance and significantly increase grain yield in greenhouse

To evaluate the grain yield under salt stress, we carefully examined the *ospqt3* mutants in the greenhouse. In the normal growth conditions the mutant plants had a little larger panicle than WT control (Figure 6A). However, in the presence of salt stress the mutant had significantly larger panicle compared to WT plants (Figure 6B). A much more significant difference was observed in the salt stress condition which, the grain yield of *ospqt3* mutant increased by more than 30 % of the wild type (Figure 6C).

**Figure 6.**
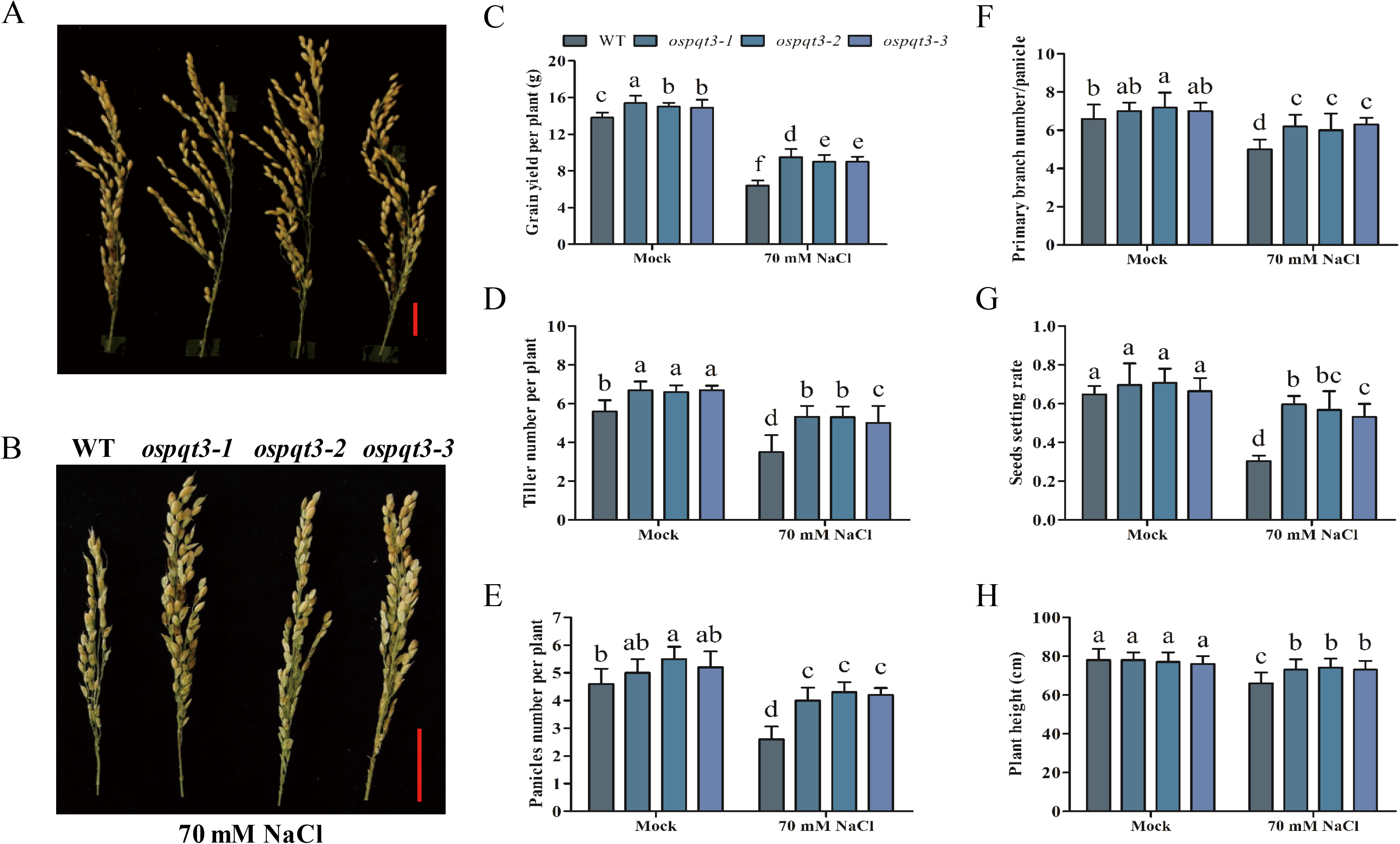
The *ospqt3* mutants exhibit enhanced yield under salt stress conditions. A and B. Representative panicles of WT, *ospqt3-1*, *ospqt3-2*, and *ospqt3-3* mutants plants grown in the greenhouse with or without 70 mM salt as described in the Methods. Bar = 4 cm. C-H. Yield components and plant height. Grain yield per plant, tillers number per plant, panicles numbers per plant, primary branch number per panicle, seeds setting rate and plant height, respectively. Values are the mean ± SD (8 plants per treatment, 3 replicates per treatment). Different letters denote significant differences (P < 0.05) from Duncan’s multiple range tests.

Tillering is one of the most vital agronomic traits related to grain production in rice (Li et al., 2003). The effective tillers number of mutant plants grown in greenhouse in the absence of salt stress increased 14 % over the WT control. While in the presence of 70 mM NaCl, the *ospqt3* mutant had significantly increased tillers number by 33.3 % more than WT plants (Figure 6D). Consistent with these results, *ospqt3* mutant grown in the field shown more tiller numbers than the wild type (Figure 4 and 5). Apparently, increased tiller number is one contributor to the increased yield. The increased panicle size of the *ospqt3* mutant rice was largely contributed by the increased number of primary branches (Figure 6E and F). The seed setting rate under well irrigated conditions shows no obvious difference between the *ospqt3* mutants and WT while in the presence of salt stress, the seeds setting rate of the *ospqt3* mutants was increased by 50 % over the WT plants (Figure 6G). Finally, *ospqt3* mutants grown in normal conditions had no clear differences in the plant height with comparison to the WT plants. However, in the salt stress conditions the plant height of the WT plants were decreased by 9 % compared to the *ospqt3* mutants (Figure 6H).

### *OsPQT3* is rapidly down-regulated by stress treatments

Quantitative PCR was used to check the expression level of *OsPQT3* in the seedling stage, vegetative and premature stages. The *OsPQT3* expression was significantly higher in the seedling shoot as compared to root. While in the vegetative and premature stages the expression was higher in root (Figure 7A). As predicted, the expression of *OsPQT3* was rapidly down-regulated upon paraquat treatment (Figure 7B), which supports that *OsPQT3* serves as negative regulator for oxidative tolerance in plants. Similarly, the expression of *OsPQT3* was down-regulated by H_2_O_2_, mannitol, drought, CdCl_2_, and salt treatment (Figure 7B-G). Relatively, paraquat and drought rapidly and significantly reduced expression of *OsPQT3* and kept the transcript level low throughout the treatments (Figure 7B and E). Although the *OsPQT3* expression was rapidly down-regulated at 0.5 and 1.0 hours of the treatment with CdCl_2_, however, *OsPQT3* transcript level was recovered at 3 and 6 hours of the treatment with CdCl_2_ (Figure 7E). These results are consistent with that of the *Arabidopsis PQT3,* indicating *PQT3* may have conserved functions in rice and Arabidopsis plants.

**Figure 7.**
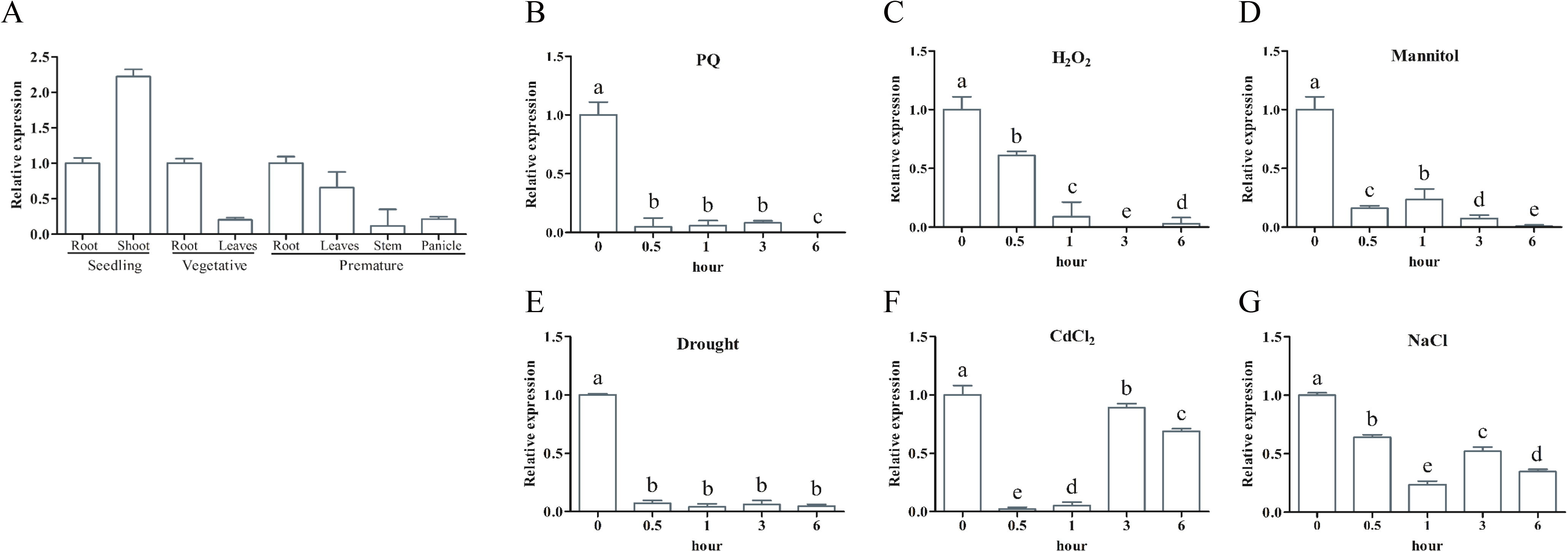
The expression pattern and rapid down-regulation of *OsPQT3* expression by various oxidative stress conditions. A. Expression pattern of *OsPQT3* in different tissues of 3 development stages using quantitative RT-PCR. *ACTIN* is used as an internal reference gene. Values are the mean ± SD (n = 3 experiments). B-G. Rapid down regulation of *OsPQT3* by stress treatment. The transcript level of *OsPQT3* was down-regulated by numerous oxidative stress: 5 μM paraquat, 10 mM H_2_O_2_, mannitol, 200 mM CdCl_2_, drought, and 150 mM NaCl. 1-week-old seedlings were used for these treatments and the RNA was extracted for quantitative RT-PCR analysis. Values are mean ± SD (n = 3 experiments). Different letters denote significant differences (P < 0.05) from Duncan’s multiple range tests.

### *OsAPX1, OsGPX1,* and *OsSOD1* are up-regulated in *ospqt3* mutants

Enzymatic defense systems are crucial for eliminating ROS. Transcript levels of glutathione peroxidase (*OsGPX*), ascorbate peroxidase (*OsAPX*), plastidic Cu/Zn *SOD* (*OsCSD2*),Cu/Zn *OsSOD* (*OsCSD1*), catalase (*OsCAT*), cytosolic mitochondrial Mn*SOD* (*OsMSD*), glutaredoxin C (*OsGRXC*), peroxiredoxin Q (*OsPRXQ*) 2-Cys peroxiredoxin B (*Os2CPB*), and atypical Cys-His rich thioredoxin (*OsACHT*) were analyzed through quantitative RT-PCR in *ospqt3* mutants and WT. We found that the transcript levels of *OsGPX1, OsAPX1* and *OsSOD1* were significantly upregulated in *ospqt3* mutants as compared to that of WT under normal conditions (Figure 8A). The up-regulated transcript levels of *OsGPX1, OsAPX1*and *OsSOD1* indicate that they are responsible for the increased resistance against oxidative stress in the mutants.

**Figure 8.**
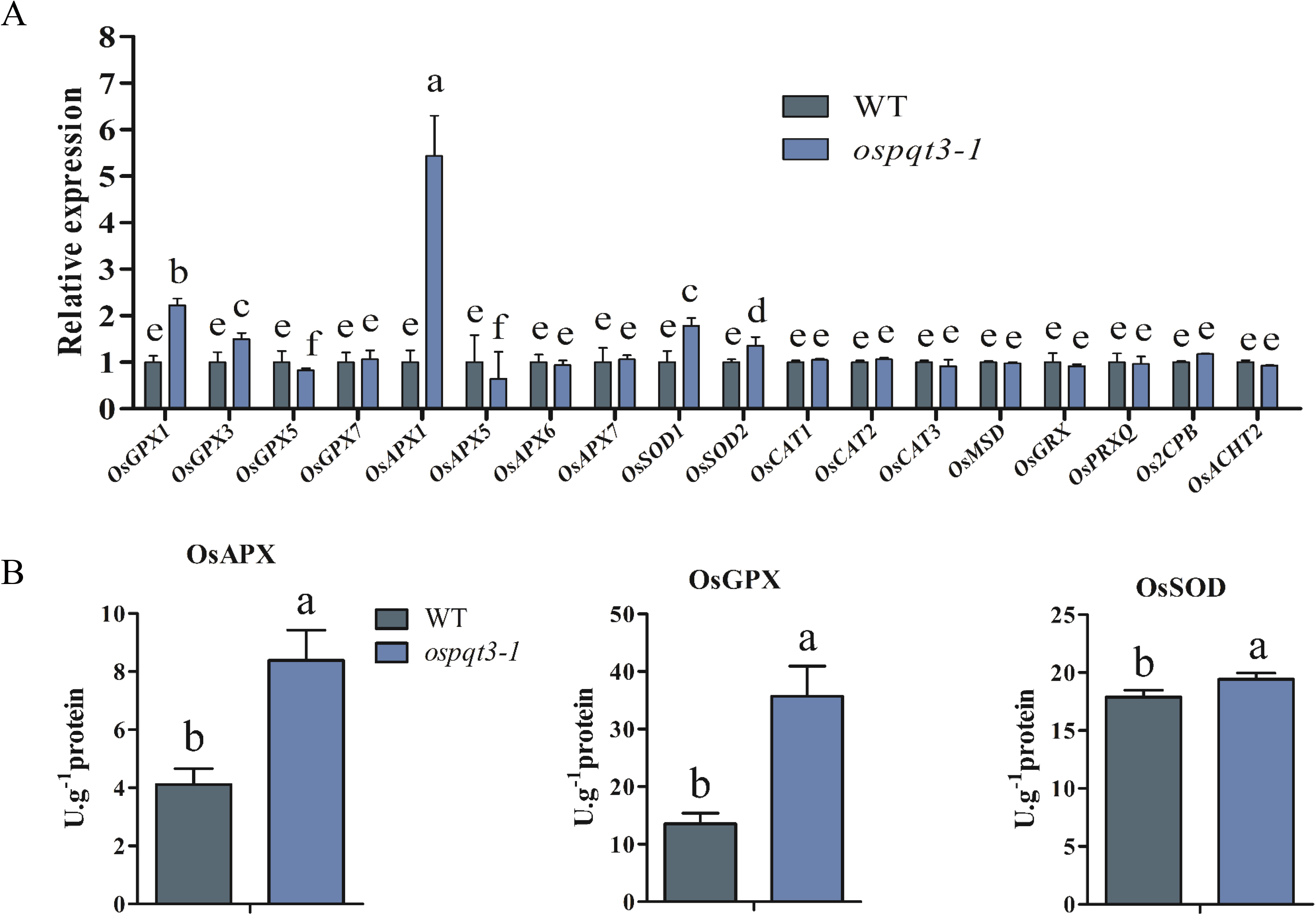
The expression of antioxidant enzymes is elevated in *pqt3* mutants. A. Quantitative RT-PCR analysis of transcript levels of antioxidant enzyme genes. RNA samples were isolated from 7-day-old WT and *ospqt3* seedlings for quantitative RT-PCR analysis. The transcript levels of *OsAPX, OsGPX, OsSOD, OsCAT, OsMSD, OsGRXC, OsPRXQ, Os2CPB*, and *OsACHT*, were analyzed. Values are mean ± SD (n = 3 experiments). Different letters denote significant differences (P < 0.05) from Duncan’s multiple range tests. B. Enzyme activity of OsAPX, OsGPX, and OsSOD, in WT and *ospqt3* mutants. Values are mean ± SD (n = 3 experiments). Different letters denote significant differences (P < 0.05) from Duncan’s multiple range tests.

We also compared the enzyme activity of APX, GPX and SOD between WT and *ospqt3* mutants. The wild type had lower enzyme activity of APX, GPX and SOD compared with *pqt3* mutants (Figure 8B). These results are consistent with that found in the Arabidopsis *pqt3* mutant, which also supports that the *PQT3* has a conserved function in rice and Arabidopsis plants. Taken together, these results demonstrate that *OsPQT3* is a negative regulator in oxidative response as *AtPQT3* in *Arabidopsis thaliana* through similar molecular mechanism.

## DISCUSSION

In this study, we assessed the function of *OsPQT3* in rice. Our results show that *ospqt3* mutants exhibit not only enhanced resistance to oxidative and salt stress but also increased grain yield of rice. These findings highlight *OsPQT3* as a promising candidate gene in crop improvement. The paraquat and salt tolerant phenotypes of *ospqt3* mutants are contributed mainly by antioxidation enzymes.

The *ospqt3* mutants showed a high seed germination rate and a high survival rate during seeding stage in the presence of paraquat and salt compared to WT plants (Figure 2). Moreover, the expression level of *OsPQT3* was significantly decreased in response to multiple different stresses such as paraquat, H_2_O_2_, mannitol, drought, CdCl_2_ and NaCl, consistent with the role of *OsPQT3* as a negative regulator. Finally, increased transcription level of *OsAPX*, *OsGPX* and *OsSOD* and increased activities of OsAPX, OsGPX and OsSOD enzymes observed in the *ospqt3* mutant plants (Figure 8) are also in agreement with their enhanced tolerance against oxidative stress. Higher concentration of ROS poses direct impact upon biological membranes, disrupts macromolecules, promotes cell senescence, induces irreversible cellular damages and eventually leads to cell death (Gill and Tuteja, 2010). Several other ROS-scavenging enzymes, in addition to *APX*, including *GPX*, *CAT*, *PrxR*, and *GST* are involved in scavenging excessive H_2_O_2_ generated under wide range of abiotic stresses (Mittler et al., 2004). These enzymes play their part in breaking H_2_O_2_ down to water and thus reduce H_2_O_2_ concentration (Dixon and Edwards, 2010; Mittler et al., 2004). It is axiomatic that up-regulating such ROS scavenger–encoding genes will speed up the process of reducing toxic ROS concentration. In *ospqt3* mutants multiple antioxidation enzymes are maintained at a higher level, which enhances the ability to resist abiotic stress. Biotic and abiotic stresses restrict crop productivity. Drought and salt are the most influential abiotic stresses that severely restrict crop productivity (Boyer, 1982; Jamil et al., 2011; Rockström and Falkenmark, 2000). The *ospqt3* mutants exhibited enhanced salt tolerance at all growth stages tested (germination, vegetative and reproductive) (Figure 2, Figure 3 and Figure 6). We observed that loss-of-*OsPQT3* affected grain yield components like effective tillers number, panicle length, primary branch number, seeds setting rate, and ultimately grain yield per plant (Figure 4 and Figure 5).

Based on our findings, *ospqt3*-enhanced grain yield in rice is primarily achieved through the following major factors. Firstly, *ospqt3* mutant plants are more tolerance to paraquat and salt due to the better protection from oxidative stress because of higher *APX*, *GPX*, and *SOD* activities (Figure 8). As an E3-ubiquitin ligase, *PQT3* plays negative role in plants to achieve oxidative tolerance via selected degradation of histone modifying PROTEIN METHYLTRANSFERASE4b (PRMT4b) which is further involved in regulating *GPX1* and *APX1* expression (Luo et al., 2016). Therefore, the *ospqt3* mutant plants performed better as indicated by their growth and yield under salt stress conditions compared with wild type plants (Figure 6). Secondly, the *ospqt3* mutant plants had more tillers and larger panicle size compared to wild type plants (Figure 4–6). The grain yield is directly connected to panicle size and effective tiller number (Xing and Zhang, 2010). The mechanism by which the *ospqt3* mutants improves the panicle size and effective tillers number is unclear yet but interesting to investigate in future.

## MATERIALS AND METHODS

### Plant material and growth conditions

Rice (*Oryza sativa*) wild type cv. Zhonghua11 along with three CRISPR/Cas edited Zhonghua11 mutants (*ospqt3-1, ospqt3-2, and ospqt3-3*) were used in this study (Hu et al., 2016). Hydroponically grown plants were maintained in Kimura B solution (EHARA et al., 1990). Medium was refreshed on alternate days while growth conditions were maintained at 28°C temperature, photo-regime of 16 hours light - 8 hours dark, 70% relative humidity, and light intensity at 200 μmol m^−2^ s^−1^.

### Seed germination

Seeds of wild type, *ospqt3* mutants were surface sterilized with (0.1 % HgCl_2_) for 15 min and then rinsed four times with sterile distilled water. To analyze seed germination, 50~70 seeds (three replicates per plant) were randomly placed in a bottle containing MS medium with paraquat or salt. All bottles were placed in a chamber that had an 8 h light (200 μmol m^−2^ s^−1^)/16 h dark photoperiod at 28°C. The seeds were considered to have germinated when their radicle or germ length reached approximately 1 mm. Seed germination was observed daily to calculate the germination percentage (Wang et al., 2010).

### Salt treatment

Wild type and mutant plants seeds, washed with distilled water, and were incubated at 37°C for 2 days. Germinated seeds were transferred to Hoagland solution (pH 6) for 3 days followed by application of 75 mM and 100 mM NaCl for 10 days. The provided growth conditions were kept at (14-h-light/10-h-dark cycle at 28°C). For the salt treatment in soil, 30 seedlings from each of the wild type, *ospqt3-1, ospqt3-2,* and *ospqt3-3* seedlings were directly sown on soil pot (the pot dimensions were 5×5×12 cm^3^, and five plants were grown per pot). After grown for 4 weeks in soil under greenhouse conditions of 16 h light/ 8 h dark) at 30°C, plants were either irrigated with 0 mM or 200 mM NaCl solution for 5-6 days.

For long term salt treatment in greenhouse, seeds of WT and *ospqt3* mutant were germinated in plates for 4 days and then sown on similar pots which used previously for salt treatment of 4-week-old seedlings. Plants were watered with 70 mM NaCl every 4-5 days, we add only water when the salt precipitation increased in the tray that holds the pots (about twice a month). The plants were grown to mature under the conditions and yield data were collected.

### Gene expression analysis

TRIZOL (Invitrogen) method was adopted for extraction of total cellular RNA while DNA was eliminated from extracted RNA through DNase I (TaKaRa). Using the cDNA synthesis kit (TaKaRa: Moloney Murine Leukemia Virus Version), cDNA was synthesized from extracted RNA. The synthesized cDNA was subjected into real-time *qPCR* using *OsACTIN1* as internal control. Primers used are enlisted in Table S1 of supplemental information.

### Enzyme activity assay

For *OsAPX*, *OsGPX*, and *OsSOD* enzyme activity assay, 16 days old rice seedlings treated hydroponically were crushed in liquid nitrogen, and re-suspended in pre-cooled enzyme extraction buffer. *In vitro* activities of *OsAPX*, *OsGPX*, and *OsSOD* were measured according to the methods described by Nakano and Asada (Nakano and Asada, 1981), Halliwell and Foyer (Halliwell and Foyer, 1978), and Mizushima et al. (Mizushima et al., 1991), respectively. Subsequently, enzyme-coupled spectrophotometer assay kit (SKBC, China) was used as per manufacturer’s guidelines to quantify the *OsAPX*, *OsGPX*, and *OsSOD* activities.

### The field trials

To test in field, plants were grown in paddy field in Hainan from December to April and in Hefei from June to August of each year. To evaluate yield components of *ospqt3* (CRISPR) mutants with Zhonghua11 background (WT) in normal field environment, three T3-homozygous independent lines of the *ospqt3* along with wild type were planted in paddy rice field, 3 plots per genotype, each plot contains 3 rows × 20 plants per row along with wild type plants. These experiments were performed in two different locations in China: Hainan Island (December to April 2019) and Hefei (April to August 2019). To reduce the variability in field test, the plants in the edge were eliminated in each plot in order to avoid margin effects.

### Evaluation of agronomic traits

Individual plant height, seed setting ratio, tiller number, number of seeds per panicle, and grain yield were measured according to protocol documented earlier (Hu et al., 2015).

### Supplemental Data

Supplemental Table S1. Primers used in this study.

### Accession Numbers

Sequence data from this article can be found in the database Rice Genome Annotation Project (https://rice.plantbiology.msu.edu/) under the following accession numbers: *OsAPX1 LOC_Os03g17690; OsPQT3 LOC_Os10g29560.1; OsAPx5, LOC_Os12g07830.1; OsAPx6, LOC_Os12g07820.1; OsAPx7,LOC_Os04g35520.2; OsGPX1,Os03g0358100; OsGPX3, LOC_Os11g18170.1; OsGPX5, LOC_Os02g44500.1; OsGPX7, LOC_Os06g08670.1; OsACHT2, LOC_Os05g11090.1; OsCAT2, LOC_Os03g45170.1; OsCAT3, LOC_Os06g51150.1; OsCSD2/ OsSOD2, LOC_Os08g44770.1; OsPRXQ, LOC_Os06g09610.1; OsGRX1, LOC_Os02g30850;* and *Os2CPB, LOC_Os04g33970.1.*

## ACKNOWLEDGEMENTS

This work was supported by the Strategic Priority Research Program of the Chinese Academy of Sciences (grant no. XDA24010303) and by grants from the National Natural Science Foundation of China (grant no. 31770273), and Ministry of Science and Technology of China (grant no. 2018ZX08009-11B, 2016ZX08005-004-003, 2016ZX08001003). Alamin Alfatih and Sami Ullah Jan are recipients of CAS-TWAS President’s Fellowship.

## AUTHOR CONTRIBUTIONS

C.B.X. and A.A. designed the experiments. A.A. performed the experiments. S.U.J., Z.S.Z., and X.J.Q. participated in field trials and data analyses. A.A. wrote the manuscript. C.B.X. and W.J. revised the manuscript. C.B.X. supervised the project.

## REFERENCES

Apel, K. and Hirt, H. (2004) Reactive oxygen species: metabolism, oxidative stress, and signal transduction. Annu. Rev. Plant Biol. 55, 373–399.

Boyer, J.S. (1982) Plant productivity and environment. Science 218, 443–448.

Chen, H., Ma, B., Zhou, Y., He, S.-J., Tang, S.-Y., Lu, X., Xie, Q., Chen, S.-Y. and Zhang, J.-S. (2018) E3 ubiquitin ligase SOR1 regulates ethylene response in rice root by modulating stability of Aux/IAA protein. Proceedings of the National Academy of Sciences 115, 4513–4518.

Chen, R.Q., Sun, S.L., Wang, C., Li, Y.S., Liang, Y., An, F.Y., Li, C., Dong, H.L., Yang, X.H., Zhang, J. and Zuo, J.R. (2009) The Arabidopsis PARAQUAT RESISTANT2 gene encodes an S-nitrosoglutathione reductase that is a key regulator of cell death. Cell Research 19, 1377–1387.

Das, K. and Roychoudhury, A. (2014) Reactive oxygen species (ROS) and response of antioxidants as ROS-scavengers during environmental stress in plants. Frontiers in Environmental Science 2, 53.

Dixon, D.P. and Edwards, R. (2010) Glutathione transferases. The Arabidopsis Book/American Society of Plant Biologists 8.

Ehara, H., Tsuchiya, M. and Ogo, T. (1990) Fundamental Growth Response to Fertilizer in Rice Plants: I. Varietal difference in the growth rate at the seedling stage. Japanese Journal of Crop Science 59, 426–434.

Gill, S.S. and Tuteja, N. (2010) Reactive oxygen species and antioxidant machinery in abiotic stress tolerance in crop plants. Plant physiology and biochemistry 48, 909–930.

Haley, T.J. (1979) Review of the toxicology of paraquat (1, 1′-dimethyl-4, 4′-bipyridinium chloride). Clinical toxicology 14, 1–46.

Halliwell, B. and Foyer, C. (1978) Properties and physiological function of a glutathione reductase purified from spinach leaves by affinity chromatography. Planta 139, 9–17.

Hu, B., Wang, W., Ou, S., Tang, J., Li, H., Che, R., Zhang, Z., Chai, X., Wang, H. and Wang, Y. (2015) Variation in NRT1. 1B contributes to nitrate-use divergence between rice subspecies. Nature Genetics 47, 834.

Hu, X., Wang, C., Fu, Y., Liu, Q., Jiao, X. and Wang, K. (2016) Expanding the range of CRISPR/Cas9 genome editing in rice. Molecular plant 9, 943–945.

Jamil, A., Riaz, S., Ashraf, M. and Foolad, M.R. (2011) Gene expression profiling of plants under salt stress. Critical Reviews in Plant Sciences 30, 435–458.

Kurepa, J., Smalle, J., Va, M., Montagu, N. and Inzé, D. (1998) Oxidative stress tolerance and longevity in Arabidopsis: the late‐flowering mutant gigantea is tolerant to paraquat. The Plant Journal 14, 759–764.

Leshem, Y., Melamed-Book, N., Cagnac, O., Ronen, G., Nishri, Y., Solomon, M., Cohen, G. and Levine, A. (2006) Suppression of Arabidopsis vesicle-SNARE expression inhibited fusion of H2O2-containing vesicles with tonoplast and increased salt tolerance. Proceedings of the National Academy of Sciences 103, 18008–18013.

Li, J.M., Su, Y.L., Gao, X.L., He, J., Liu, S.S. and Wang, X.W. (2011) Molecular characterization and oxidative stress response of an intracellular Cu/Zn superoxide dismutase (CuZnSOD) of the whitefly, Bemisia tabaci. Archives of insect biochemistry and physiology 77, 118–133.

Li, X., Qian, Q., Fu, Z., Wang, Y., Xiong, G., Zeng, D., Wang, X., Liu, X., Teng, S. and Hiroshi, F. (2003) Control of tillering in rice. Nature 422, 618.

Lim, S.D., Cho, H.Y., Park, Y.C., Ham, D.J., Lee, J.K. and Jang, C.S. (2013) The rice RING finger E3 ligase, OsHCI1, drives nuclear export of multiple substrate proteins and its heterogeneous overexpression enhances acquired thermotolerance. Journal of experimental botany 64, 2899–2914.

Luo, C., Cai, X.-T., Du, J., Zhao, T.-L., Wang, P.-F., Zhao, P.-X., Liu, R., Xie, Q., Cao, X.-F. and Xiang, C.-B. (2016) PARAQUAT TOLERANCE3 is an E3 ligase that switches off activated oxidative response by targeting histone-modifying PROTEIN METHYLTRANSFERASE4b. PLoS genetics 12, e1006332.

Marino, D., Froidure, S., Canonne, J., Khaled, S.B., Khafif, M., Pouzet, C., Jauneau, A., Roby, D. and Rivas, S. (2013) Arabidopsis ubiquitin ligase MIEL1 mediates degradation of the transcription factor MYB30 weakening plant defence. Nature communications 4, 1476.

Miller, G., Suzuki, N., Ciftci‐Yilmaz, S. and Mittler, R. (2010) Reactive oxygen species homeostasis and signalling during drought and salinity stresses. Plant, cell & environment 33, 453–467.

Mittler, R., Vanderauwera, S., Gollery, M. and Van Breusegem, F. (2004) Reactive oxygen gene network of plants. Trends in plant science 9, 490–498.

Mizushima, Y., Hoshi, K., Yanagawa, A. and Takano, K. (1991) Topical application of superoxide dismutase cream. Drugs under experimental and clinical research 17, 127–131.

Moon, J., Parry, G. and Estelle, M. (2004) The ubiquitin-proteasome pathway and plant development. The Plant Cell 16, 3181–3195.

Nakano, Y. and Asada, K. (1981) Hydrogen peroxide is scavenged by ascorbate-specific peroxidase in spinach chloroplasts. Plant and cell physiology 22, 867–880.

Rockström, J. and Falkenmark, M. (2000) Semiarid crop production from a hydrological perspective: gap between potential and actual yields. Critical reviews in plant sciences 19, 319–346.

Summers, L.A. (1980) The bipyridinium herbicides:Academic Press Inc.

Takahashi, M. and Asada, K. (1988) Superoxide production in aprotic interior of chloroplast thylakoids. Archives of biochemistry and biophysics 267, 714–722.

Van Breusegem, F., Bailey-Serres, J. and Mittler, R. (2008) Unraveling the tapestry of networks involving reactive oxygen species in plants. Plant physiology 147, 978–984.

Vierstra, R.D. (2009) The ubiquitin–26S proteasome system at the nexus of plant biology. Nature Reviews Molecular Cell Biology 10, 385.

Wang, Z.-f., Wang, J.-f., Bao, Y.-m., Wang, F.-h. and Zhang, H.-s. (2010) Quantitative trait loci analysis for rice seed vigor during the germination stage. Journal of Zhejiang University Science B 11, 958–964.

Xi, J., Xu, P. and Xiang, C.B. (2012) Loss of AtPDR11, a plasma membrane-localized ABC transporter, confers paraquat tolerance in Arabidopsis thaliana. Plant Journal 69, 782–791.

Xing, Y. and Zhang, Q. (2010) Genetic and Molecular Bases of Rice Yield. Annual Review of Plant Biology 61, 421–442.

Ye, B. and Gressel, J. (2000) Transient, oxidant-induced antioxidant transcript and enzyme levels correlate with greater oxidant-resistance in paraquat-resistant Conyza bonariensis. Planta 211, 50–61.

Yu, Y., Xu, W., Wang, J., Wang, L., Yao, W., Yang, Y., Xu, Y., Ma, F., Du, Y. and Wang, Y. (2013) The Chinese wild grapevine (Vitis pseudoreticulata) E3 ubiquitin ligase Erysiphe necator‐induced RING finger protein 1 (EIRP1) activates plant defense responses by inducing proteolysis of the VpWRKY11 transcription factor. New Phytologist 200, 834–846.

Zhou, S., Chen, Q., Sun, Y. and Li, Y. (2017) Histone H2B monoubiquitination regulates salt stress‐induced microtubule depolymerization in Arabidopsis. Plant, cell & environment 40, 1512–1530.

