## supplemental Table S1 for "Loss of rice *PARAQUAT TOLERANCE 3* confers enhanced resistance to abiotic stresses and increases grain yield in field"

| *Primers* | Sequences |
| --- | --- |
| *OsPQT3-CRISPR-Cas9-G1-F* | CTGAGTGGGCATTATGGCTG |
| *OsPQT3-CRISPR-Cas9-G1-R* | CTCAGTTATCCACAACAATCCT |
| *OsPQT3-CRISPR-Cas9-G2-F* | CGGTTGTTGAGTGCATGTATC |
| *OsPQT3-CRISPR-Cas9-G2-R* | GGCATGGTTTCTTCTACCCTA |
| *qPCR-OsPQT3-F* | TACAAGTTCAAGAGTGCCAGAG |
| *qPCR-OsPQT3-R* | ATGGTTTCTTCTACCCTACCTT |
| *OsAPx1-F* | ATCAAGGAGGAGATACCCACCAT |
| *OsAPx1-R* | TGGTCAGAACCCTTGGTAGCAT |
| *OsAPx5-F* | ATGGCCGTCGTGCACCGCAT |
| *OsAPx5-R* | CTAACCGAACCAGGATGGGAT |
| *OsAPx6-F* | AGTCCACCTCCTGCCATCCCA |
| *OsAPx6-R* | CATTCACAAGACCTGCATTAGC |
| *OsAPx7-F* | CTCCTCCACCTCCTCCGCCT |
| *OsAPx7-R* | AGTGCGTCGTCTTGAGGAGCT |
| *OsGPX1-F* | TGCTAATCGTCGTCAACGTCG |
| *OsGPX1-R* | GCTGGTCAGATCCTGGCTCTT |
| *OsGPX3-F* | ATGGCGGCTACAACTACAAGTA |
| *OsGPX3-R* | GTGGACGGAGGTGGGCAGGT |
| *OsGPX5-F* | TTCATCTCTCGCCTCCGGCT |
| *OsGPX5-R* | GGACGGACCAGCGGCGAGA |
| *OsGPX6-F* | TGGGCTCTGGTATTGACGACG |
| *OsGPX6-R* | GCAGAACTCTACCACTGCTTC |
| *OsGPX7-F* | ATGGCGTCCACCACCACCAC |
| *OsGPX7-R* | GAAGTCGTGGACGCTCTTCC |
| *OsCAT1-F* | AGCTTGCACAGTTTGACAGGGA |
| *OsCAT1-R* | GTTCGGTTCTCCACAGTCGTG |
| *OsCAT2-F* | TCGTTCAACGGCCCGCTGTG |
| *OsCAT2-R* | GATACGCTCCCTGTCGAAGTT |
| *OsCAT3-F* | AGCTTGCACAGTTTGACAGGGA |
| *OsCAT3-R* | ACGACTGTGGAGAACCGAACA |
| *OsACHT2-F* | TGCTCCTCTGCCTCCTCCTCC |
| *OsACHT2-R* | GTCTGAGACGGCTGCATGGG |
| *OsSOD2-F* | ATGCAAGCCATCCTCGCCGC |
| *OsSOD2-R* | TTGAGCACGGCGACGGCCTT |
| *OsGRX-F* | TGGTGGTGTTCAGCGTGAGCA |
| *OsGRX-R* | GCTTGCCGCCGATGAAGACGA |
| *OsPRXQ-F* | ATGGCATTCGCGGTCTCCAC |
| *OsPRXQ-R* | CAGACGATCCTGTTCCTCCC |
| *Os2CPB-F* | GATCACTGCTTTCAGCGACAG |
| *Os2CPB-R* | GAATGCTGAATCACACCCTCC |

**Table S1. Primers used in this study.**
